# Glucolipotoxicity-induced Mig6 desensitizes EGFR signaling and promotes pancreatic beta cell death

**DOI:** 10.1101/2020.11.24.380279

**Authors:** Yi-Chun Chen, Andrew J. Lutkewitte, Halesha D. Basavarajappa, Patrick T. Fueger

## Abstract

A loss of functional beta cell mass is a final etiological event in the development of frank type 2 diabetes (T2D). To preserve or expand beta cells and therefore treat/prevent T2D, growth factors have been considered therapeutically but have largely failed to achieve robust clinical success. The molecular mechanisms preventing the activation of mitogenic signaling pathways from maintaining functional beta cell mass during the development of T2D remain unknown. We speculated that endogenous negative effectors of mitogenic signaling cascades impede beta cell survival/expansion. Thus, we tested the hypothesis that a stress-inducible epidermal growth factor receptor (EGFR) inhibitor, Mitogen-inducible gene 6 (Mig6), regulates beta cell fate in a T2D milieu. To this end, we determined that: 1) glucolipotoxicity (GLT) induces Mig6, thereby blunting EGFR signaling cascades, and 2) Mig6 mediates molecular events regulating beta cell survival/death. We discovered that GLT impairs EGFR activation, and Mig6 is elevated in human islets from T2D donors as well as GLT-treated rodent islets and 832/13 INS-1 beta cells. Mig6 is essential for GLT-induced EGFR desensitization, as Mig6 suppression rescued the GLT-impaired EGFR and ERK1/2 activation. Further, Mig6 mediated EGFR but not insulin-like growth factor-1 receptor nor hepatocyte growth factor receptor activity in beta cells. Finally, we identified that elevated Mig6 augmented beta cell apoptosis, as Mig6 suppression reduced apoptosis during GLT. In conclusion, we established that T2D and GLT induce Mig6 in pancreatic beta cells. The elevated Mig6 desensitizes EGFR signaling and induces beta cell death. Our findings suggest that Mig6 could be a novel therapeutic target for T2D, as blocking Mig6 could possibly enhance mitogenic signaling cascades in a diabetic milieu to promote beta cell survival and prevent beta cell death.

---

In normal physiological conditions, to ensure proper, robust insulin release, beta cell insulin secretory capacity (i.e., beta cell function) and insulin stores (dependent on beta cell mass) are tightly regulated to match demand. For example, during pregnancy and obesity, functional beta cell mass (the composite of beta cell mass and functional capacity) is increased to maintain optimal glycemic control. Studies have suggested that growth factor receptor signaling pathways, such as epidermal growth factor receptor (EGFR) signaling, are among the primary contributors of beta cell mass expansion (14, 15) by promoting beta cell replication and survival. Unfortunately, the capacity to increase, and even preserve, functional beta cell mass is finite.

The inability of beta cells to meet a sustained demand to produce and secrete insulin imposed by systemic insulin resistance precipitates type 2 diabetes (T2D), where de-compensation of functional beta cell mass occurs. In the period of pre-diabetes leading up to frank T2D, bouts of hyperglycemia and persistent hyperlipidemia (glucotoxicity and lipotoxicity, respectively, and hereafter jointly referred to as glucolipotoxicity, or GLT) accentuate beta cell failure. Interestingly, the activation of proximal signaling molecules of growth factor receptors (e.g., Akt and glycogen synthase kinase 3/GSK3) is inhibited in beta cells under stressed conditions (7, 30, 32). Such observations suggest that activation of the growth factor receptors themselves might be inhibited during the development of T2D, thereby limiting beta cell proliferation and survival. Thus, we hypothesized that the glucolipotoxic milieu central to the development of T2D suppresses the activation of beta cell growth factor receptors, thereby inactivating survival signaling cascades, and contributing to beta cell loss through apoptosis. Further, we speculate that endogenous feedback inhibition is a mechanism for suppression of growth factor signaling in beta cells in T2D.

In many cells, EGFR signaling is engaged to promote cellular replication and survival; indeed, EGFR signaling is central to preserving beta cells *in vivo* and in *vitro*. Mice expressing constitutively-active EGFR in the pancreas are protected against the development of beta cell toxin-induced diabetes, in that their beta cell loss is inhibited (13). Conversely, pharmacologically blocking Raf-1 signaling downstream of EGFR induces beta cell death (1). As alluded to above, EGFR signaling is also required for beta cell expansion, as mice expressing a dominant negative form of EGFR fail to acquire compensatory beta cell expansion during pregnancy and obesity (14, 15). Thus, inactivation of EGFR signaling has dire consequences for functional beta cell mass. Nevertheless, little attention has been given to the molecular mechanisms controlling inactivation of growth factor signaling in the context of T2D.

Like most signaling cascades, activation of a pathway is followed by inactivation through classical negative feedback; EGFR signaling is no different. We have been investigating the impact of endogenous feedback inhibition of EGFR by the adapter protein Mig6 on functional beta cell mass (5, 6). Here, we report that GLT induces Mig6 where it terminates EGFR signaling and promotes beta cell apoptosis. Thus, suppressing the actions of Mig6 could be fruitful for re-engaging pro-survival signaling in beta cells in T2D.

## MATERIALS AND METHODS

### Cell culture, reagents, and the use of adenovirus

INS-1-derived 832/13 and 828/33 rat insulinoma cells were grown in 11.1 mM D-glucose RPMI-1640 medium supplemented with 10% fetal bovine serum, 100 U/ml penicillin, 100 µg/ml streptomycin, 10 mmol/l HEPES, 2 mmol/l L-glutamine, 1 mmol/l sodium pyruvate, and 50 µmol/l β-mercaptoethanol, as preciously described (16, 31). 832/13 cells are glucose-responsive for secreting insulin and sensitive to apoptosis. 828/33 cells stably overexpress Bcl-2 and are thus resistant to apoptosis.

In lipotoxicity experiments, sodium palmitate (Sigma; St. Louis, MO) was dissolved in 0.1M NaOH buffer at 70°C, and mixed with 5% fatty acid-free BSA (Sigma) solution at 37°C to yield a 5 mM palmitic acid-BSA complex stock solution. Palmitic acid-BSA complex was diluted into serum free RPMI culture medium to obtain various concentrations of palmitic acid ranging from 0.1 to 0.4 mM. In glucotoxicity experiments, 25 mM glucose, serum-free RPMI culture medium supplemented with 0.1% BSA was used as high glucose treatment, and 5 mM glucose plus 20 mM D-mannitol (osmotic control; Sigma) was used as low glucose control. For glucolipotoxicity experiments, 0.4 mM palmitic acid plus 25 mM glucose medium was used to create glucolipotoxicity.

For EGF stimulation experiments, 832/13 cells were challenged with glucolipotoxicity for 4 h, starved in RPMI 1640 medium containing 2.5 mM glucose and 0.1% BSA for 2 h, and treated with 10 ng/ml rat recombinant EGF (R&D Systems) for 5 min. For growth factors stimulation experiments, 832/13 cells were starved for 2 h, and treated with insulin-like growth factor 1 (IGF-1, NIH repository) or hepatocyte growth factor (HGF, R&D Systems) for various times.

In gene overexpression studies, recombinant adenoviral vectors expressing Mig6 or green fluorescent protein (GFP) under the control of cytomegalovirus (CMV) promoter were used as previously descried (6). In gene suppression studies, recombinant adenoviral vectors expressing small interfering RNAs (siRNA) specific to rat Mig6 or with no know gene homology (scrambled siRNA) were used as previously described (3). Alternatively, for cell apoptosis experiments, Mig6 siRNA (Mig6 ON-TARGET plus Smartpool, Dharmacon) was transfected with Lipofectamine 2000 (Invitrogen) to suppress Mig6 expression, and non-targeting siRNA served as a negative control. Cells were challenged by glucolipotoxicity 72 h after transfection, and apoptosis assays were performed.

### Human and rodent islet experiments

Cadaveric human islets were obtained from the Integrated Islet Distribution Program or the Southern California Islet Resource Center affiliated with the Beckman Research Institute of the City of Hope. Islets from four donors with T2D and four donors without diabetes were analyzed to determine *MIG6* expression levels in mRNA isolated from islets upon receipt to the laboratory (typically one day after isolation). In signaling experiments, human islets from eight donors with BMIs lower than 30 were cultured similar to cell experiments with the exception that CMRL-1066 media with 5 mM glucose and supplemented with 10% fetal bovin serum, 50 units/ml penicillin, and 50 µg/ml streptomycin was used.

Rat pancreatic islets were collected from male Wistar rats weighing approximately 250 g (25, 26). After collagenase digestion, islets were hand-picked and cultured in 5 mM glucose RPMI medium supplemented with 10% fetal bovin serum, 50 units/ml penicillin, and 50 µg/ml streptomycin overnight before drug treatments.

### Apoptosis assays

Following siRNA treatment (48 h), 40,000 832/13 cells were plated in black-walled, 96-well plates. The following day, media was replaced with starvation media, and cells were treated with either BSA or palmitic acid-coupled BSA for times indicated. Caspase 3/7 activity was measured using an Apolive-Glo kit (Promega, Madison, WI) and measured using a SpectraMax M5 (Molecular Devices, Sunnyvale, CA). Briefly, following GLT treatments cells were incubated with Caspase-Glo 3/7 Reagent for 30 mins at RT and luminescence was measured.

### Immunoblot analysis

Cells were lysed in 1% IGEPAL reagent supplemented with 10% glycerol, 16 mM NaCl, 25 mM HEPES, 60 mM n-octylglucoside (Research Products International Corp.), phosphatase inhibitor cocktails (PhosSTOP tablets, Roche), and protease inhibitor cocktails (EDTA-free cOmplete tablets, Roche). Lysates were resolved on a 10% NuPAGE Bis-Tris Gel (Invitrogen), transferred to an Immobilon-FL Transfer Membrane (Millipore), and incubated with primary antibodies (**Table 1**). Subsequently, membranes were incubated with IRDye 800 or 700 fluorophore-labeled secondary antibodies from LI-COR. Protein bands were visualized using the Odyssey System (LI-COR) and quantified with Image J software (NIH).

**Table 1.** List of antibodies used for immunoblotting.

| **Name** | **Vendor, model number** | **Dilution** |
| --- | --- | --- |
| Anti-Actin | MP Biomedicals, #691002 | 1:5000 |
| Anti-Akt | Cell Signaling, #2920 | 1:1000 |
| Anti-caspase 3 | Cell Signaling, #9662 | 1:1000 |
| Anti-CHOP | Santa Cruz, #7351 | 1:250 |
| Anti-EGFR | Sigma-Aldrich, #E3138 | 1:1000 |
| Anti-eIF2α | Cell Signaling, #5324 | 1:1000 |
| Anti-ERK1/2 | Cell Signaling, #4696 | 1:1000 |
| Anti-γ-tubulin | Sigma-Aldrich, #T6557 | 1:5000 |
| Anti-Gapdh | Abcam, #Ab9483 | 1:5000 |
| Anti-Mig6 | Santa Cruz, #D-1 | 1:250 |
| Anti-phospho-Akt (Thr308) | Cell Signaling, #4056 | 1:1000 |
| Anti-phospho-EGFR (Tyr1068) | Cell Signaling, #3777 | 1:1250 |
| Anti-phospho-eIF2α (Ser51) | Cell Signaling, #3398 | 1:1000 |
| Anti-phospho-ERK1/2 (Thr202/Tyr204) | Cell Signaling, #4370 | 1:2000 |
| IRDye 800 or 700 fluorophore-conjugated antibodies | LI-COR | 1:10000 |

Phosphorylated protein levels were normalized to the total protein levels in the cell lysate, and the total (e.g., non-phosphorylated) protein levels were normalized to tubulin or GAPDH protein levels.

### Quantitative RT-PCR analysis

RNA from 832/13 cells, rat islets, and human islets were isolated using RNeasy Mini or Micro kits (Qiagen; Valencia, CA). Reverse transcription was completed with a High Capacity cDNA Reverse Transcription kit (Applied Biosystems). The threshold cycle methodology was used to calculate the relative quantities of the mRNA products of *Mig6, Socs4, Socs5, Frs3*, and *Lrig1* (**Table 2**). PCR reactions were performed in triplicate for each sample from at least three independent experiments and were normalized to *Gapdh* or *beta-actin* gene expression levels.

**Table 2.**
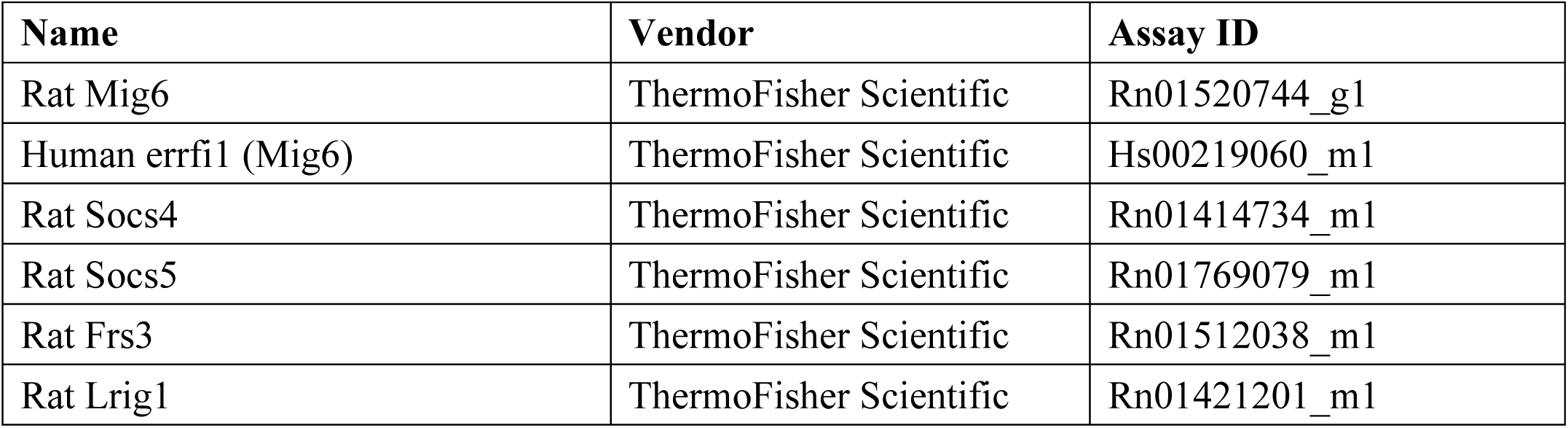
List of Taqman gene expression assays.

### Statistical analysis

All data are reported as means ± SEM. Protein and mRNA data were normalized to control conditions and were presented as relative expression. Student’s *t*-test or ANOVA (with Bonferroni *post-hoc* tests) were performed using GraphPad Prism software to detect statistical differences (*P* < 0.05).

## RESULTS

### GLT and ER stress attenuate EGFR activation in rodent beta cells and human islets

To study the extent to which GLT compromises EGFR activation, we exposed human islets and 832/13 INS-1 cells to medium containing high glucose and palmitic acid and assessed phosphorylation of EGFR. We identified that GLT treatment prevents EGF-mediated EGFR phosphorylation in both human islets and the beta cell line (**Figure 1**). Notably, neither the basal phosphorylation nor total cellular abundance of EGFR was changed by GLT (data not shown), indicating that the attenuated EGFR phosphorylation and activation were likely the consequences of EGFR kinase interruption. Because GLT imposes endoplasmic reticulum (ER) stress on beta cells and triggers deleterious effects such as beta cell death, indicated by phosphorylation of eIF2α and JNK as well as elevated cleaved caspase 3 (**Figures 1A and 2A-B**), we examined the extent to which ER stress induction alone was sufficient to attenuate EGFR activation. Pretreatment with thapsigargin (the sarcoendoplasmic reticulum Ca(2+) ATPase 2 pump inhibitor) induces ER stress and inhibited EGFR activation in 832/13 cells (**Figure 2C-D**). The above findings suggest that the pathological stress stimuli present in T2D compromise the activation of EGFR. Importantly, cell death in GLT *per se* does not decrease EGFR phosphorylation, as beta cells overexpressing Bcl-2, which confers resistance to apoptosis, were still sensitive to GLT with respect to its ability to prevent maximal EGFR phosphorylation (**Figure 3**).

**Figure 1.**
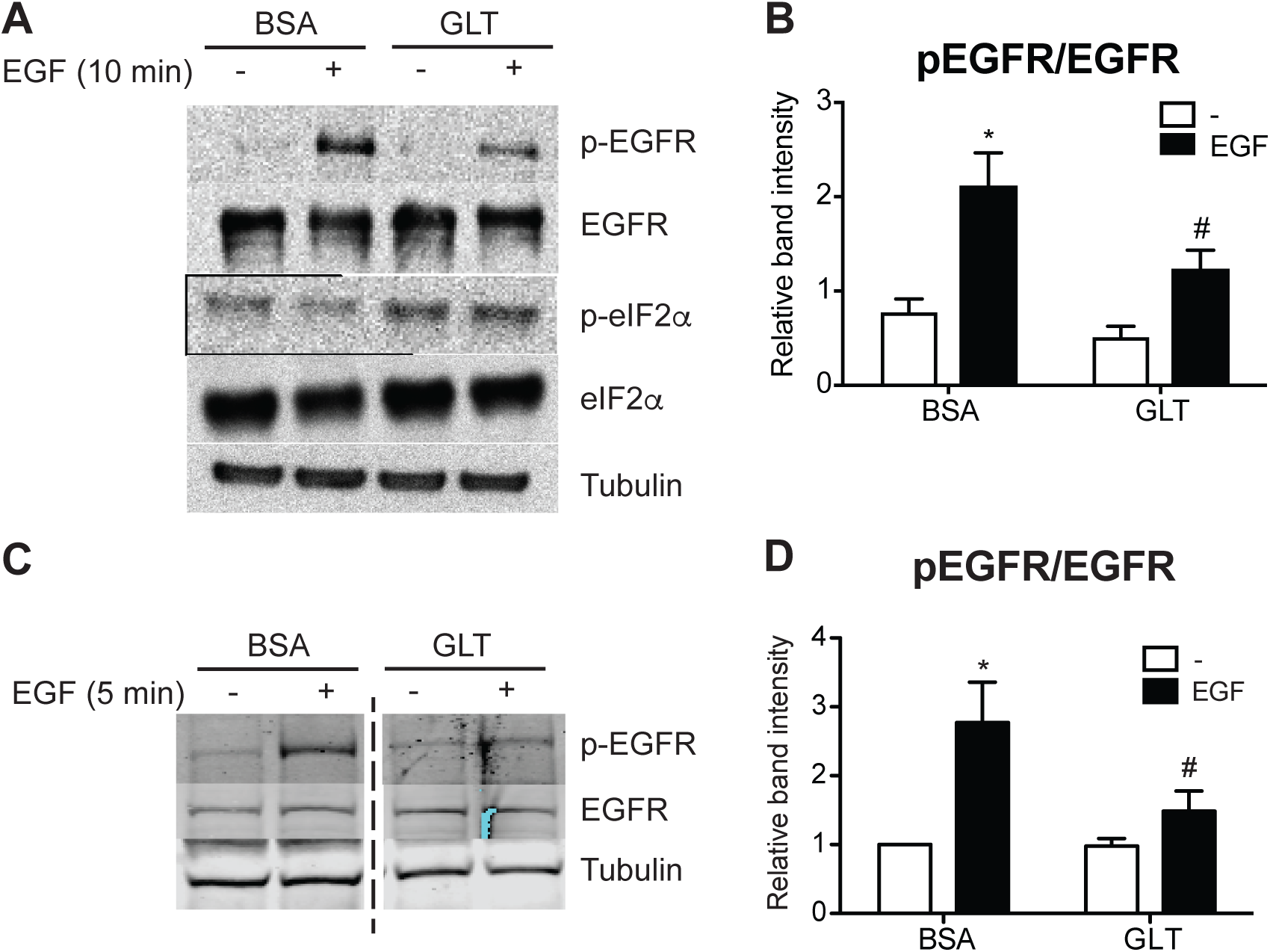
Glucolipotoxicity impairs EGFR activation in beta cells and human islets. Human islets and 832/13 cells were cultured in media with 5 mM glucose and BSA, or 25 mM glucose and 400 µM palmitic acid (glucolipotoxicity, GLT) for 48 or 8 h, respectively, followed by starvation in 5 mM glucose medium for 2 h, and then stimulated with recombinant human EGF (50 ng/ml for 10 min for islets and 10 ng/ml for 5 min for cells). Protein levels of p-EGFR, EGFR, p-eIF2α, and tubulin were analyzed by immunoblotting. Shown are representative immunoblots from human islets (**A-B)** and 832/13 cells (**C-D**), and quantified data are reported as fold induction relative to BSA, non-stimulated samples. *n* = 8 independent human islet preparations and 3 independent cell line experiments; * *p* < 0.05 vs. BSA, non-stimulated; # *p* < 0.05 vs. BSA, EGF-stimulated.

**Figure 2.**
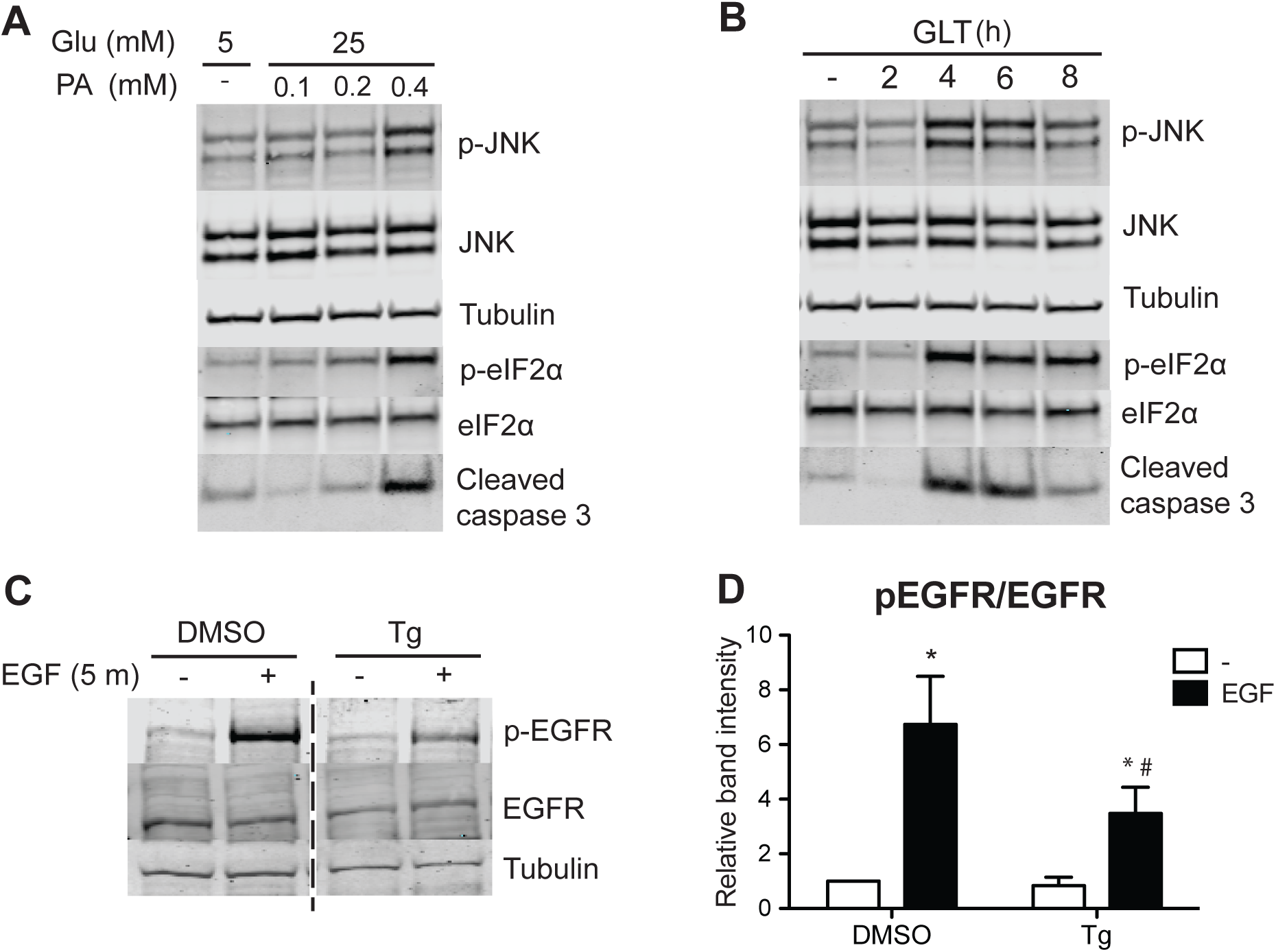
ER stress impairs EGFR phosphorylation. **(A)** To verify induction of ER stress, 832/13 cells were cultured in media containing 5 mM glucose and BSA or 25 mM glucose and increasing concentrations of palmitic acid (0, 0.1, 0.2, or 0.4 mM) for 8 h. **(B)** In a complementary experiment, cells were exposed to 25 mM glucose and 0.4 mM palmitic acid for up to 8 h. Cells were harvested and lysates were immunoblotted using antibodies directed against p-JNK, JNK, p-eIF2α, eIF2α, cleaved caspase 3 to establish the extent of ER stress produced by glucolipotoxicity. Shown are representative, confirmatory immunoblots. **(C)** To induce ER stress, 832/13 cells were treated with DMSO or 1 µM thapsigargin (Tg) for 4 h, followed by starvation and EGF stimulation as before. Protein levels of p-EGFR, EGFR, and tubulin were analyzed by immunoblotting. Representative blots of n ≥ 3 experiments are shown and results quantified in **(D)**. \**p* < 0.05 vs. BSA, non-stimulated; #*p* < 0.05 vs. BSA, EGF-stimulated.)

**Figure 3.**
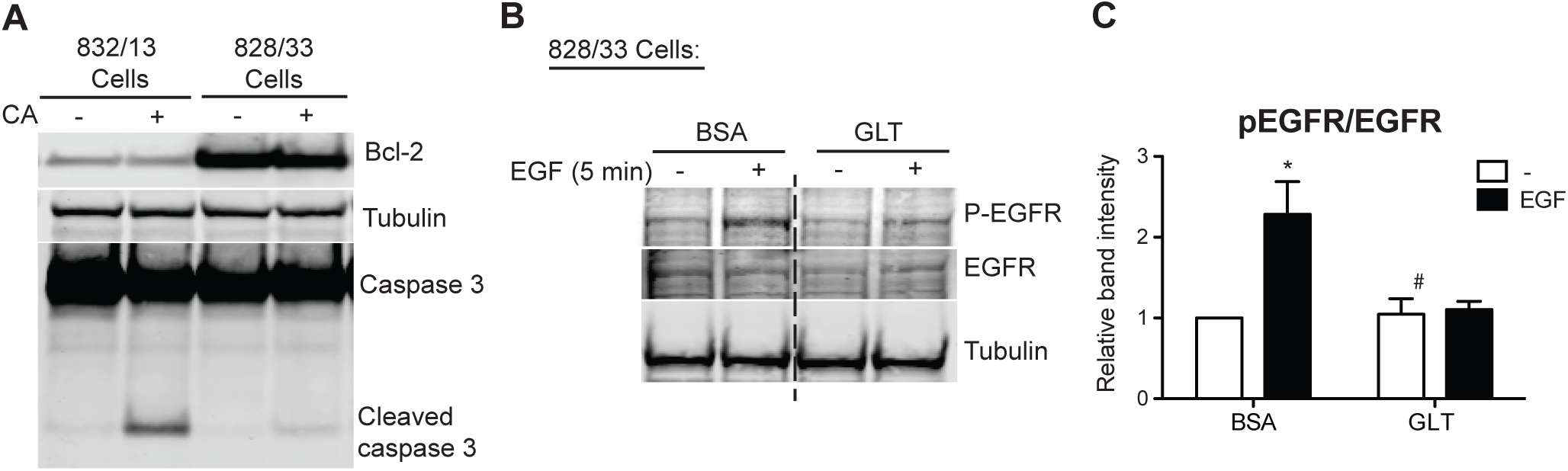
Apoptosis is not required for glucolipotoxicity-impaired EGFR activation. **(A)** To promote apoptosis, 832/13 and Bcl-2 overexpressing 828/33 cells were treated with 1 µM camptothecin (CA) for 8 h. To demonstrate resistance to cell death in 828/33 cells, protein levels of Bcl-2, full-length and cleaved caspase 3, and tubulin were analyzed by immunoblotting. **(B)** 828/33 cells were cultured in media as in Figure 1. Protein levels of p-EGFR, EGFR, and tubulin were analyzed by immunoblotting, and quantified results are reported in **(C)**. *n* = 3; \**p* < 0.05 vs. all other groups.

### EGFR feedback inhibitor Mig6 is elevated in GLT-treated beta cells and T2D human islets

To identify the factors associated with EGFR inactivation during GLT, we examined a set of well-defined, inducible EGFR inhibitors (29). First, we discovered that Mig6, an adaptor protein that blocks EGFR activation, was induced by GLT whereas expression of other EGFR inhibitors -- SOCS4, SOCS5, FRS3, and LRIG1 -- all remained unchanged by GLT (**Figure 4**). To further study the specific effects of glucotoxicity and lipotoxicity on the induction of Mig6, we treated the 832/13 cells with high levels of glucose and/or palmitic acid and measured Mig6 expression levels (**Figure 5**). We established that high glucose induced Mig6 in a dose-dependent manner that was not due to the osmotic actions of glucose (**Figure 5A-B**). However, palmitic acid alone did not induce Mig6 at the concentrations and time points examined (**Figure 5C-D**). In addition, because high glucose stimulates insulin secretion in the beta cells, we needed to determine the extent to which the autocrine or paracrine effects of insulin might promote Mig6 expression; exogenous insulin treatment did not alter Mig6 expression levels (**Figure 5E**). Finally, we identified that Mig6 mRNA expression was greater in islets isolated from donors with T2D compared to control donors, and Mig6 mRNA and protein expression was induced in rodent islets exposed to GLT (**Figure 6**). These data suggested that diabetogenic stress induces Mig6 in beta cells.

**Figure 4.**
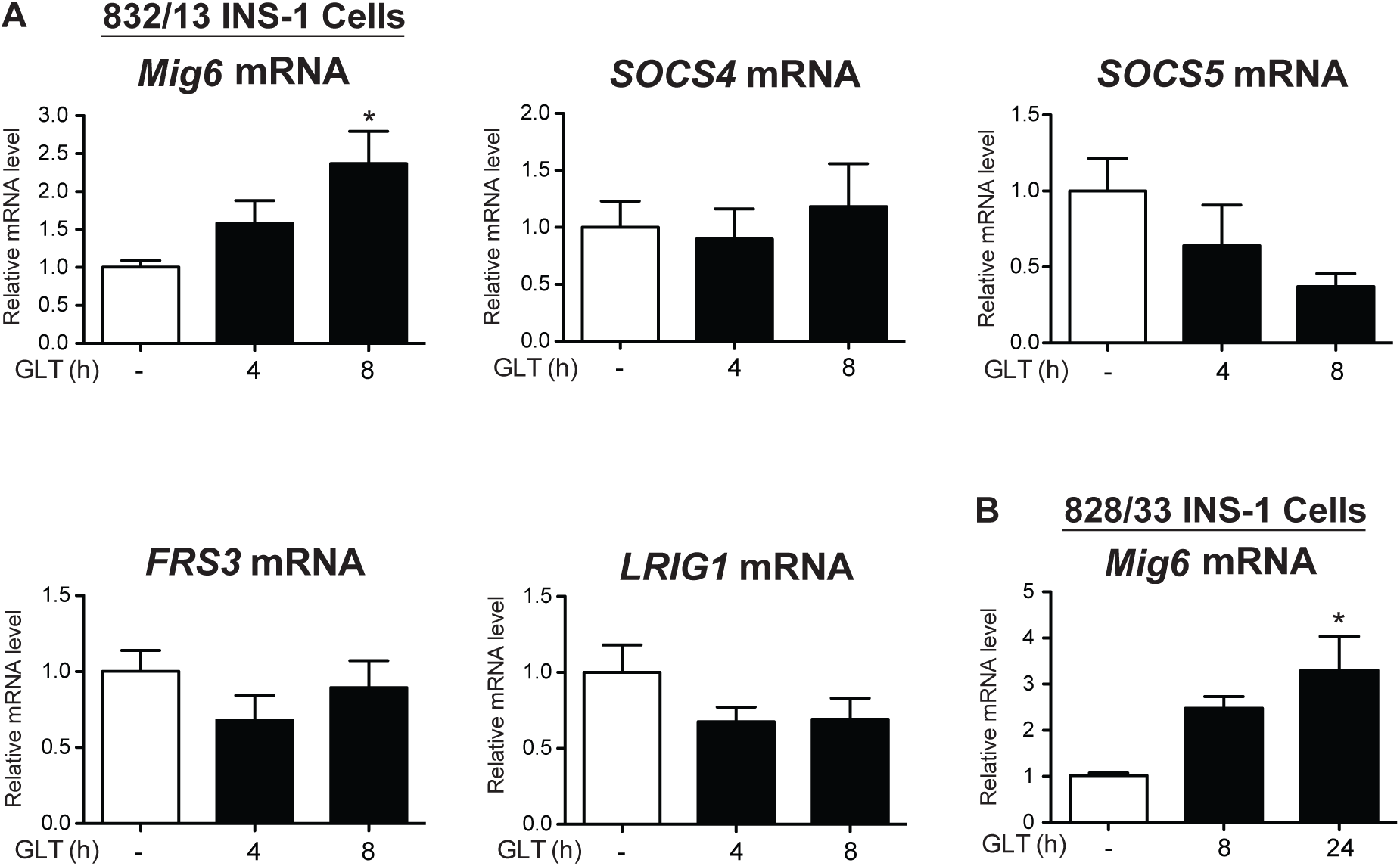
Mig6, but not other feedback inhibitors of EGFR, is induced by glucolipotoxicity. **(A)** 832/13 and **(B)** 828/33 cells were cultured in control (white bars) or glucolipotoxic (black bars) media for up to 8 h. Expression of *Mig6, SOCS4, SOCS5, FRS1*, and *LRIG1* mRNA was quantified using qRT-PCR. *n* = 3; \**p* < 0.05 vs. control media.

**Figure 5.**
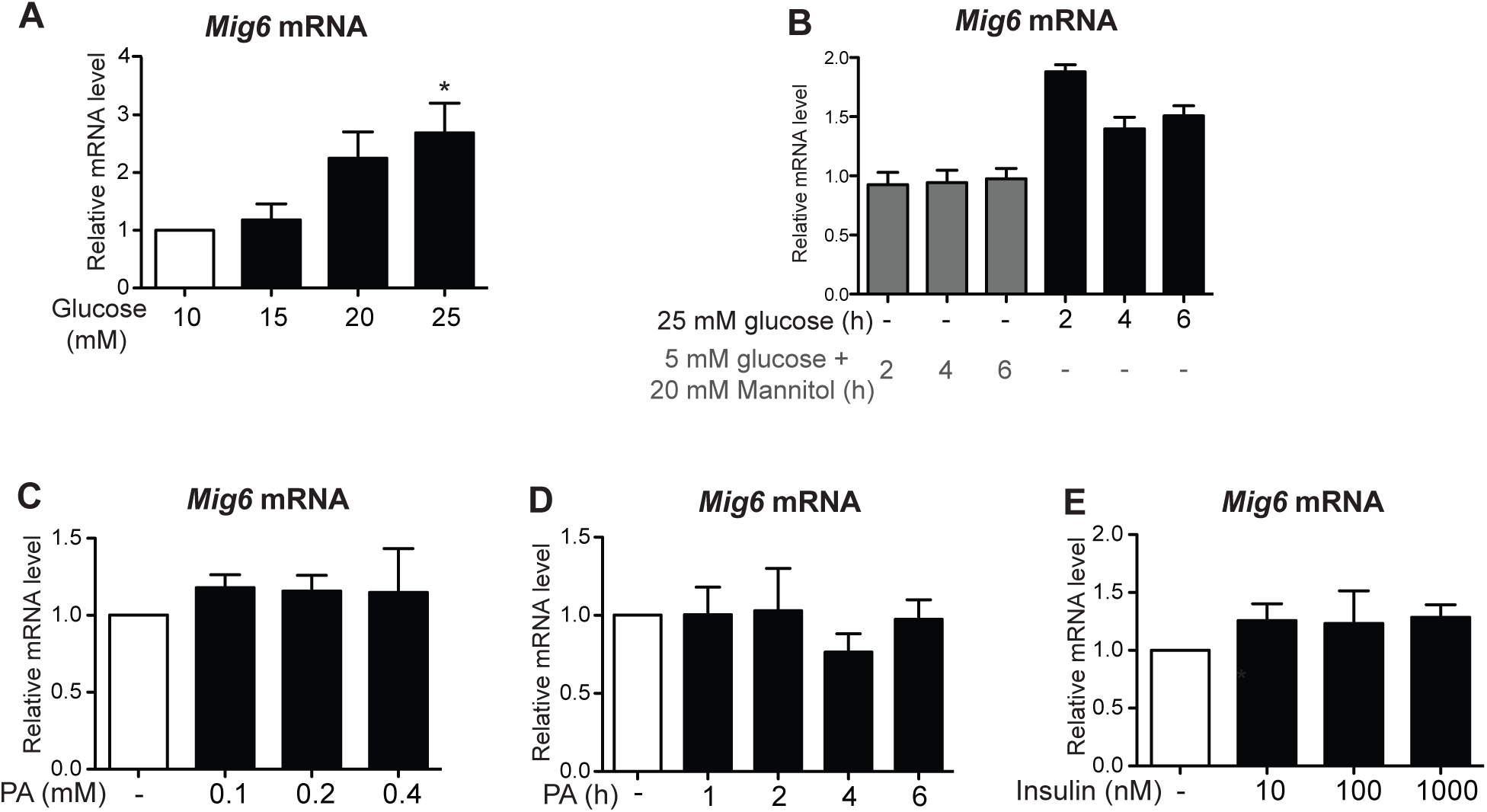
Gluco-, but not lipotoxicity, alone induces Mig6 expression. 832/13 cells were treated with (A) 5, 10, 15, 20, or 25 mM glucose for 4 h, **(B)** 25 mM glucose or 5 mM glucose + 20 mM mannitol (as an osmotic stress control) for 0, 2, 4, or 6 h, **(C)** BSA, 100, 200, 400 µM palmitic acid for 4 h, **(D)** 400 µM palmitic acid for the indicated times, or **(E)** 0, 10, 100, or 1000 nM recombinant human insulin for 4 h. *Mig6* mRNA levels were determined by qRT-PCR. n ≥ 3 experiments. \**p* < 0.05 vs. BSA / 5 mM glucose + 20 mM mannitol.

**Figure 6.**
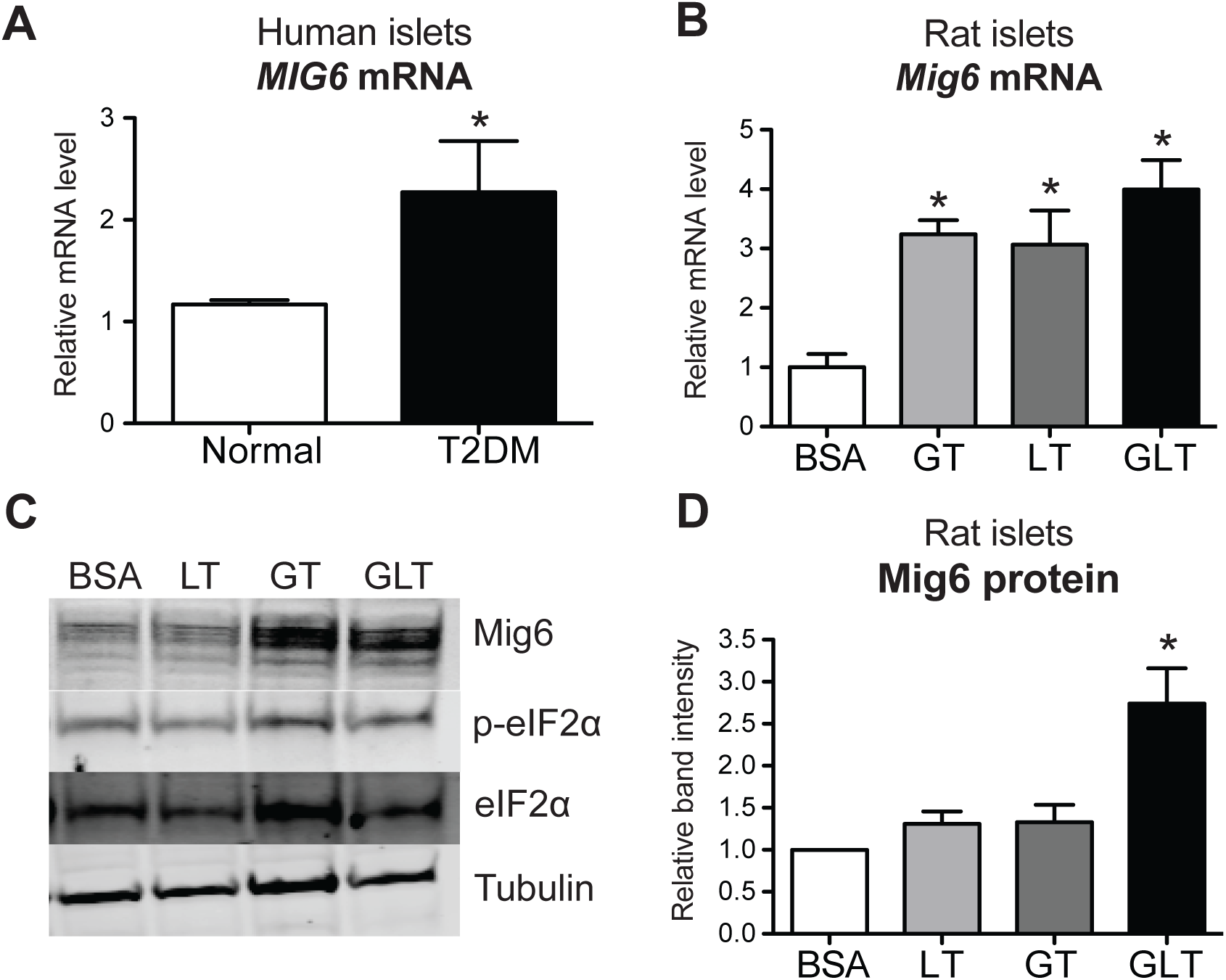
Mig6 is elevated in T2DM human and rodent islets treated with glucolipotoxicity. **(A)** *MIG6* mRNA was measured in human islets from normal (white bar) and type 2 diabetic (black bar) cadaver donors. **(B)** Rat islets were cultured in media containing 5 mM glucose and BSA (white bar), 25 mM glucose and BSA (glucotoxicity, GT; light gray bar), 5 mM glucose and 0.4 mM palmitic acid (lipotoxicity, LT; dark gray bar), or 25 mM glucose and 0.4 mM palmitic acid (glucolipotoxicity, GLT; black bar) for 8 h. *Mig6* mRNA was measured by RT-PCR. **(C)** Rat islet lysates were immunoblotted with antibodies directed against Mig6, p-eIF2α, eIF2α, and tubulin, and **(D)** results for Mig6 content were quantified. *n* = 3-4; \**p* < 0.05 vs. normal or 5 mM glucose with BSA.

### Mig6 inhibits EGFR in pancreatic beta cells and promotes death during GLT

Because GLT hinders EGFR activation and the inhibitor of EGFR, Mig6, is induced by GLT, it is intuitive to speculate that stress-inducible Mig6 controls EGFR inactivation during GLT. Thus, we used a RNA interference approach to examine the functional significance of Mig6 in EGFR signaling. Suppression of Mig6 enhanced both EGFR and ERK1/2 phosphorylation following EGF stimulation (**Figure 7A-C**). In contrast, elevated Mig6 expression dampened downstream ERK1/2 phosphorylation (**Figure 7D-F**). Importantly, the actions of Mig6 were restricted to EGFR signaling, as altering Mig6 expression (overexpression or suppression) did not alter hepatocyte growth factor-stimulated ERK1/2 phosphorylation or insulin-like growth factor-1-stimulated Akt phosphorylation (**Figure 7G-H**).

**Figure 7.**
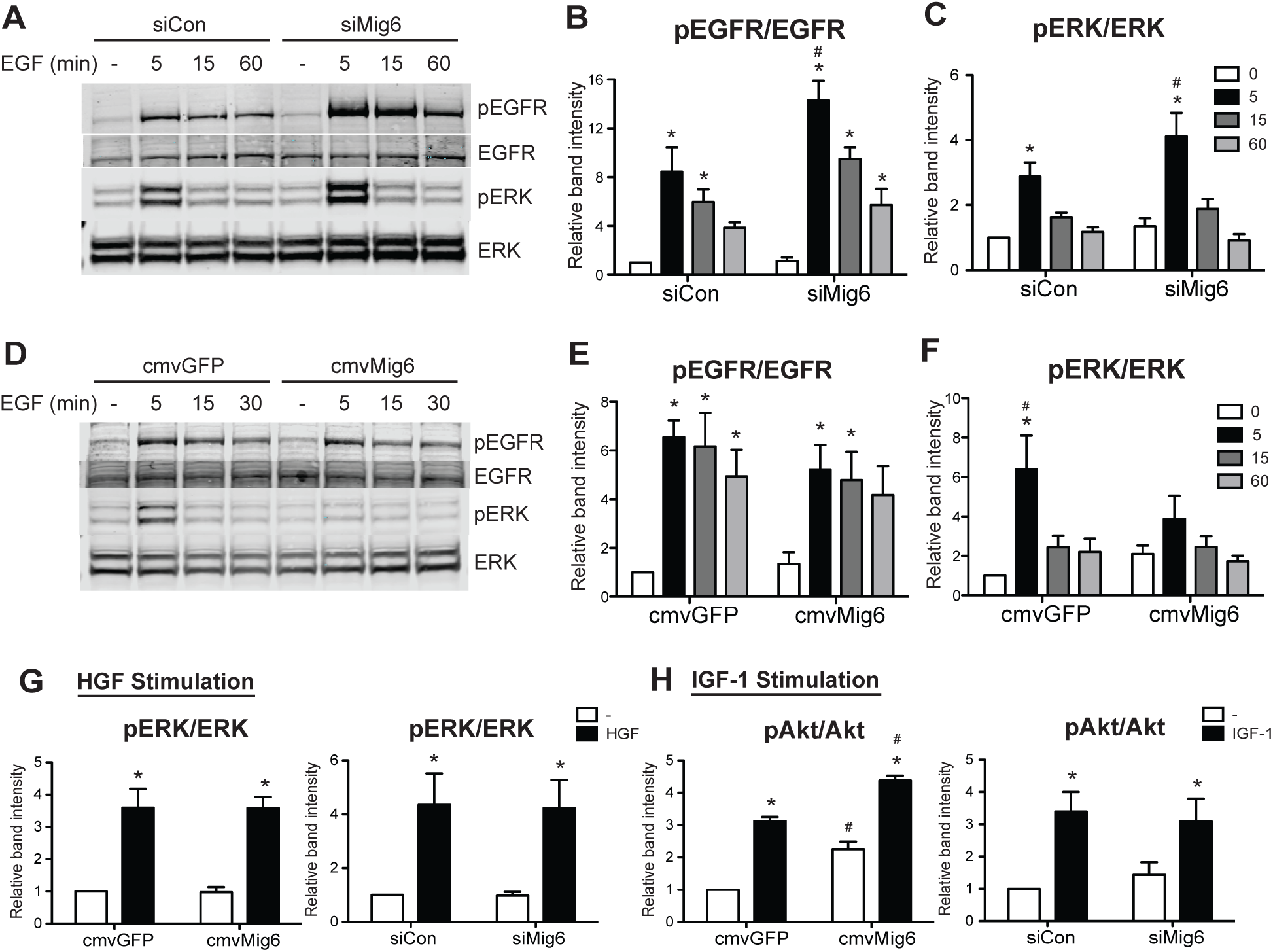
Mig6 controls EGF, but neither IGF1 nor HGF, pro-survival signaling pathways. **(A-F)** 832/13 cells were transduced with adenoviruses carrying cmvGFP vs. cmvMig6 or siCon vs. siMig6. Post transduction, cells were starved in 5mM glucose and 0.1% BSA medium for 2 h, followed by 10 ng/ml recombinant rat EGF stimulation for 5 min. **(G, H)** After adenoviral transduction and starvation as in A & D, 832/13 cells were treated with recombinant human HGF or IGF-1 for 5 min. Protein levels of p-EGFR, p-Erk, Erk, p-Akt, Akt, and tubulin were analyzed by immunoblotting. n ≥ 3. \**p* < 0.05 vs. non-stimulated. ^#^*p* < 0.05 vs. control virus, stimulated condition.

Finally, as stress-induced Mig6 suppresses the EGFR signaling pathway, we sought to determine the extent to which silencing Mig6 would restore EGFR activity and prevent beta cell death during GLT. Again, using siRNA-mediated suppression of Mig6 (**Figure 8A**), we determined that reducing Mig6 increased EGFR and ERK1/2 phosphorylation during GLT (**Figure 8B-D**). Importantly, Mig6 suppression limited beta cell apoptosis in GLT, as measured by caspase 3/7 activity (**Figure 8E**).

**Figure 8.**
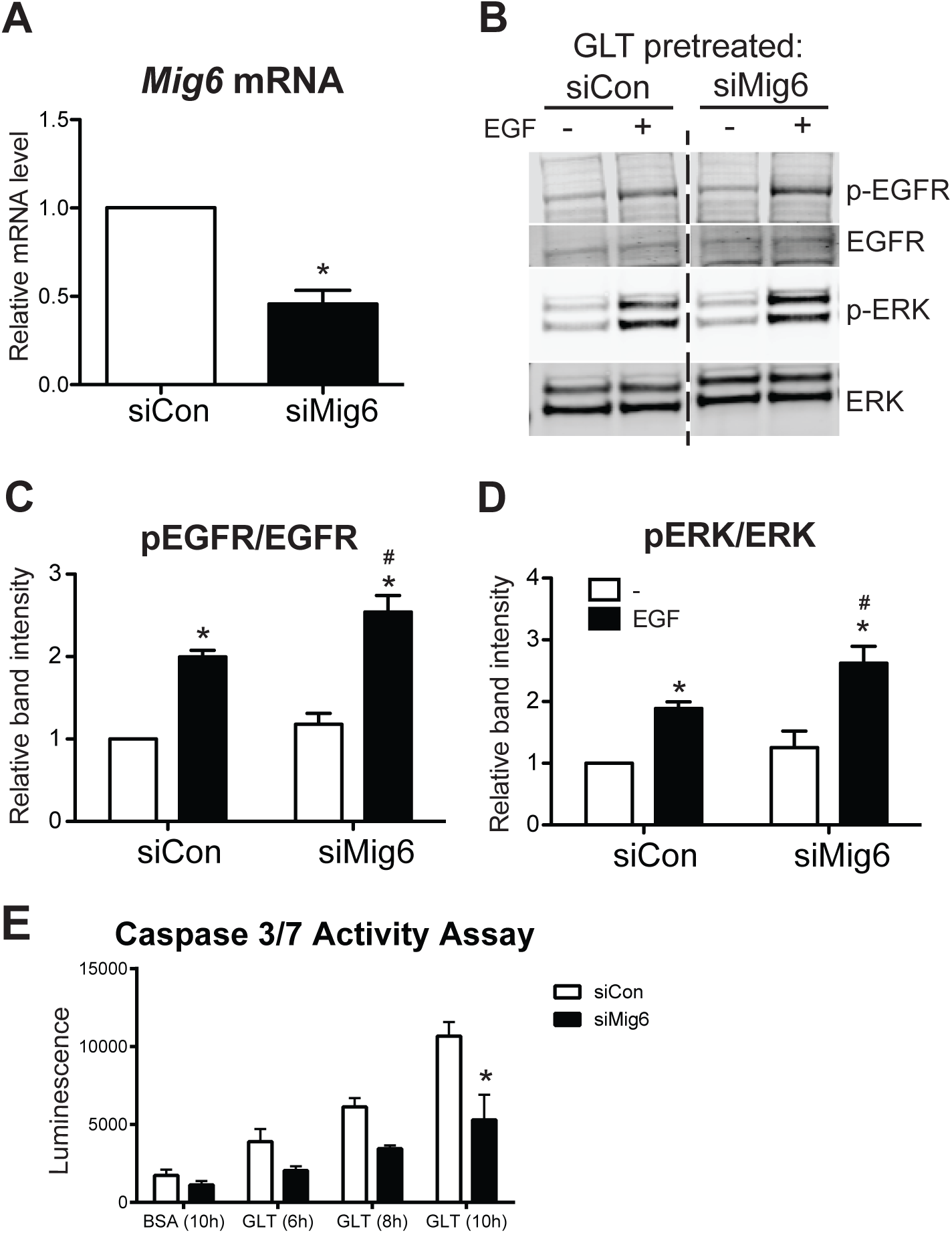
Mig6 suppression dampens apoptosis during glucolipotoxicity. **(A)** 832/13 cells were transduced with adenoviral vectors carrying either a scrambled control siRNA (siCon) or shRNA sequence against Mig6 (siMig6). *Mig6* mRNA levels were determined by qRT-PCR. *N* = 4; \**p* < 0.05. **(B-D)** Transduced cells were treated with GLT and EGF as described previously. Protein levels of p-EGFR, EGFR, p-Erk, and Erk were determined by immunoblotting. Data are reported as fold induction related to the GLT-treated, non-EGF-stimulated group. n ≥ 3. \**p* < 0.05 vs. EGF-treated. ^#^*p* < 0.05 vs. siCon EGF-stimulated. **(E)** Caspase 3/7 activity was measured following exposure to glucolipotoxic conditions in 832/13 cells siCon or siMig6. *n* = 3 experiments; \**p* < 0.05 vs. siCon.

## DISCUSSION

Although genetic manipulation of pancreatic EGFR in mice leads to the acceleration or prevention of diabetes, the natural history of EGFR kinase activity in different phases during the progression of diabetes remain unknown (13-15, 24). In regards to beta cell de-compensation, the phase of declining functional beta cell mass prior to the onset of frank T2D, the extent to which diabetogenic stress stimuli alter EGFR activity and impact beta cell life/death decisions is unclear. It has been reported that diabetic stressors could compromise the activation and propagation of receptor tyrosine kinase (RTK) signaling cascades in pancreatic beta cells (12, 20). For example, GLT and cytokine challenges hinder the activation of insulin receptor and downstream phosphatidylinositol 3-kinase, hence preventing the cyto-protective effects of insulin in beta cells (7, 8, 32). However, the molecular mechanisms responsible for this stress-mediated RTK inactivation remain to be defined. It is likely that there are stress-responsive factors that crosstalk with RTK signaling machinery in pathological conditions.

In this study, we demonstrated that glucolipotoxicity and ER stress attenuate EGFR activation in pancreatic beta cells via the stress-responsive EGFR inhibitor, Mig6. Mig6 was initially characterized as an endogenous EGFR feedback inhibitor but has also been suggested to impair other RTKs (2, 27, 33); yet in our work here, we have been unable to ascribe actions of Mig6 to HGF or IGF-1 signaling in 832/13 rat pancreatic beta cells. After mitogen stimulation, Mig6 is activated to abolish EGFR signaling transmission via a two-tiered mechanism: 1) Mig6 binds to the EGFR intracellular kinase domain and inhibits kinase dimerization and activation, and 2) Mig6 facilitates EGFR endo-lysosomal sorting and degradation (9). However, a new role of Mig6 as a stress-induced modulator cellular signaling and function has also been revealed. Makkinje at el. first reported that mechanical stress in diabetic nephropathy is sufficient to induce Mig6, and the transient expression of Mig6 results in selective activation of JNK (23). Later, Mabuchi et al. further suggested that Mig6 is able to bind to I kappa B alpha, resulting in NF-kappa B activation (21). Additionally, Hopkins et al. demonstrated that ligand deprivation promotes Mig6-mediated c-Abl activation and cell death (17). Recent work has suggested that Mig6 modulates the DNA damage response by interacting with the serine/threonine kinase ATM and histone H2AX (19). Furthermore, as previously reported by our group, both ER stress and pro-inflammatory cytokines both induce Mig6, and haploinsuficiency of Mig6 prevents mice from developing an experimentally-induced form of T1D (4, 5).

Here, we established that GLT-induced beta cell apoptosis is mediated, at least in part, by the induction and actions of Mig6. The deleterious Mig6-mediated effects could be EGFR dependent and/or independent. Beyond the well-described feedback inhibition of EGFR signaling, Mig6 activates pro-apoptotic JNK via its Cdc42/Rac interactive binding domain, representing an EGFR independent response (22). In addition to Mig6, there are likely other stress-mediated factors controlling EGFR inactivation and downstream alterations in ERK1/2 signaling. For instance, cell surface EGFR could be modified and inhibited by advanced-glycation precursors present in GLT, and cellular stress-activated phosphatases could also inactivate EGFR (10, 18, 28). Importantly, we established that several other feedback inhibitors of EGFR – Socs4, Socs5, Frs3, and Lrig1 – were not induced by GLT in beta cells, thus highlighting the importance of Mig6 in EGFR feedback inhibition. ERK1/2 signaling can also be modulated by factors others than EGFR in the context of GLT, and thus pathways beyond EGFR-Mig6 must be considered. Nevertheless, we have provided evidence that the feedback inhibition of EGFR in GLT is likely mediated by the direct actions of Mig6.

This work presents a potentially novel mechanism for reduced beta cell proliferation and survival in the states of chronic over-nutrition. It is known that short-term intralipid infusion enhances beta cell proliferation via EGFR and mTOR signaling pathways in adult rodents (34), but chronic nutrient overload (i.e., high-fat diet feeding) does not promote beta cell mass expansion (11). The contribution of Mig6 to restraining beta cell proliferation and survival during obesity *in vivo* remains to be determined. However, as Mig6 is elevated in islets isolated from T2D patients, we speculated that Mig6 perhaps contributes to the dampening of EGFR signaling activation during the progression of T2D.

In summary, we discovered that GLT attenuates EGFR activation via Mig6; and Mig6 modulates GLT-induced beta cell apoptosis. This work highlights the broad effects that a diabetogenic milieu has on cellular signaling and suggests that reactivation of pro-survival RTK signaling could be beneficial for fortifying functional beta cell mass. How Mig6 might control beta cell survival beyond the direct feedback inhibition of EGFR remains to be determined. Thus, we propose that Mig6 might be a suitable therapeutic target to promote beta cell survival in preventing and treating T2D.

## Funding

P.T.F. was supported by grants and funding from the National Institutes of Health (DK078732 and DK099311). Y.-C.C. was supported by a DeVault Fellowship from Indiana University School of Medicine. A.J.L. was supported by a Predoctoral Fellowship from the American Heart Association Midwest Region.

## Contributions

Yi-Chun Chen conceived the experimental design, performed experiments, analyzed data, and wrote and edited the manuscript; Andrew J. Lutkewitte performed experiments, analyzed data, and edited the manuscript; Halesha D. Basavarajappa performed experiments, analyzed data, and edited the manuscript; and Patrick T. Fueger conceived the experimental design, analyzed data, and wrote and edited the manuscript.

## Disclosures

The authors have nothing to disclose.

## Notes

### Competing Interest Statement

The authors have declared no competing interest.

## BIBLIOGRAPHY

1. Alejandro EU, and Johnson JD. Inhibition of Raf-1 alters multiple downstream pathways to induce pancreatic beta-cell apoptosis. The Journal of biological chemistry 283: 2407–2417, 2008.

2. Anastasi S, Fiorentino L, Fiorini M, Fraioli R, Sala G, Castellani L, Alema S, Alimandi M, and Segatto O. Feedback inhibition by RALT controls signal output by the ErbB network. Oncogene 22: 4221–4234, 2003.

3. Bain JR, Schisler JC, Takeuchi K, Newgard CB, and Becker TC. An adenovirus vector for efficient RNA interference-mediated suppression of target genes in insulinoma cells and pancreatic islets of langerhans. Diabetes 53: 2190–2194, 2004.

4. Chen YC, Colvin ES, Griffin KE, Maier BF, and Fueger PT. Mig6 haploinsufficiency protects mice against streptozotocin-induced diabetes. Diabetologia 2014.

5. Chen YC, Colvin ES, Maier BF, Mirmira RG, and Fueger PT. Mitogen-inducible gene 6 triggers apoptosis and exacerbates ER stress-induced beta-cell death. Molecular endocrinology 27: 162–171, 2013.

6. Colvin ES, Ma HY, Chen YC, Hernandez AM, and Fueger PT. Glucocorticoid-induced suppression of beta-cell proliferation is mediated by Mig6. Endocrinology 154: 1039–1046, 2013.

7. Cousin SP, Hugl SR, Wrede CE, Kajio H, Myers MG, Jr., and Rhodes CJ. Free fatty acid-induced inhibition of glucose and insulin-like growth factor I-induced deoxyribonucleic acid synthesis in the pancreatic beta-cell line INS-1. Endocrinology 142: 229–240, 2001.

8. Fornoni A, Pileggi A, Molano RD, Sanabria NY, Tejada T, Gonzalez-Quintana J, Ichii H, Inverardi L, Ricordi C, and Pastori RL. Inhibition of c-jun N terminal kinase (JNK) improves functional beta cell mass in human islets and leads to AKT and glycogen synthase kinase-3 (GSK-3) phosphorylation. Diabetologia 51: 298–308, 2008.

9. Frosi Y, Anastasi S, Ballaro C, Varsano G, Castellani L, Maspero E, Polo S, Alema S, and Segatto O. A two-tiered mechanism of EGFR inhibition by RALT/MIG6 via kinase suppression and receptor degradation. The Journal of cell biology 189: 557–571, 2010.

10. Goldkorn T, Ravid T, and Khan EM. Life and death decisions: ceramide generation and EGF receptor trafficking are modulated by oxidative stress. Antioxidants & redox signaling 7: 119–128, 2005.

11. Golson ML, Misfeldt AA, Kopsombut UG, Petersen CP, and Gannon M. High Fat Diet Regulation of beta-Cell Proliferation and beta-Cell Mass. The open endocrinology journal 4: 2010.

12. Grempler R, Leicht S, Kischel I, Eickelmann P, and Redemann N. Inhibition of SH2-domain containing inositol phosphatase 2 (SHIP2) in insulin producing INS1E cells improves insulin signal transduction and induces proliferation. FEBS letters 581: 5885–5890, 2007.

13. Hakonen E, Ustinov J, Eizirik DL, Sariola H, Miettinen PJ, and Otonkoski T. In vivo activation of the PI3K-Akt pathway in mouse beta cells by the EGFR mutation L858R protects against diabetes. Diabetologia 57: 970–979, 2014.

14. Hakonen E, Ustinov J, Mathijs I, Palgi J, Bouwens L, Miettinen PJ, and Otonkoski T. Epidermal growth factor (EGF)-receptor signalling is needed for murine beta cell mass expansion in response to high-fat diet and pregnancy but not after pancreatic duct ligation. Diabetologia 54: 1735–1743, 2011.

15. Hakonen E, Ustinov J, Palgi J, Miettinen PJ, and Otonkoski T. EGFR signaling promotes beta-cell proliferation and survivin expression during pregnancy. PloS one 9: e93651, 2014.

16. Hohmeier HE, Mulder H, Chen G, Henkel-Rieger R, Prentki M, and Newgard CB. Isolation of INS-1-derived cell lines with robust ATP-sensitive K+ channel-dependent and - independent glucose-stimulated insulin secretion. Diabetes 49: 424–430, 2000.

17. Hopkins S, Linderoth E, Hantschel O, Suarez-Henriques P, Pilia G, Kendrick H, Smalley MJ, Superti-Furga G, and Ferby I. Mig6 is a sensor of EGF receptor inactivation that directly activates c-Abl to induce apoptosis during epithelial homeostasis. Developmental cell 23: 547–559, 2012.

18. Knebel A, Rahmsdorf HJ, Ullrich A, and Herrlich P. Dephosphorylation of receptor tyrosine kinases as target of regulation by radiation, oxidants or alkylating agents. The EMBO journal 15: 5314–5325, 1996.

19. Li C, Park S, Zhang X, Eisenberg LM, Zhao H, Darzynkiewicz Z, and Xu D. Nuclear Gene 33/Mig6 regulates the DNA damage response through an ATM serine/threonine kinase-dependent mechanism. The Journal of biological chemistry 292: 16746–16759, 2017.

20. Lu B, Wu H, Gu P, Du H, Shao J, Wang J, and Zou D. Improved glucose-stimulated insulin secretion by intra-islet inhibition of protein-tyrosine phosphatase 1B expression in rats fed a high-fat diet. Journal of endocrinological investigation 35: 63–70, 2012.

21. Mabuchi R, Sasazuki T, and Shirasawa S. Mapping of the critical region of mitogene-inducible gene-6 for NF-kappaB activation. Oncology reports 13: 473–476, 2005.

22. Makkinje A, Quinn DA, Chen A, Cadilla CL, Force T, Bonventre JV, and Kyriakis JM. Gene 33/Mig-6, a transcriptionally inducible adapter protein that binds GTP-Cdc42 and activates SAPK/JNK. A potential marker transcript for chronic pathologic conditions, such as diabetic nephropathy. Possible role in the response to persistent stress. J Biol Chem 275: 17838–17847, 2000.

23. Makkinje A, Quinn DA, Chen A, Cadilla CL, Force T, Bonventre JV, and Kyriakis JM. Gene 33/Mig-6, a transcriptionally inducible adapter protein that binds GTP-Cdc42 and activates SAPK/JNK. A potential marker transcript for chronic pathologic conditions, such as diabetic nephropathy. Possible role in the response to persistent stress. The Journal of biological chemistry 275: 17838–17847, 2000.

24. Miettinen PJ, Ustinov J, Ormio P, Gao R, Palgi J, Hakonen E, Juntti-Berggren L, Berggren PO, and Otonkoski T. Downregulation of EGF receptor signaling in pancreatic islets causes diabetes due to impaired postnatal beta-cell growth. Diabetes 55: 3299–3308, 2006.

25. Milburn JL, Jr., Hirose H, Lee YH, Nagasawa Y, Ogawa A, Ohneda M, BeltrandelRio H, Newgard CB, Johnson JH, and Unger RH. Pancreatic beta-cells in obesity. Evidence for induction of functional, morphologic, and metabolic abnormalities by increased long chain fatty acids. The Journal of biological chemistry 270: 1295–1299, 1995.

26. Naber SP, McDonald JM, Jarett L, McDaniel ML, Ludvigsen CW, and Lacy PE. Preliminary characterization of calcium binding in islet-cell plasma membranes. Diabetologia 19: 439–444, 1980.

27. Pante G, Thompson J, Lamballe F, Iwata T, Ferby I, Barr FA, Davies AM, Maina F, and Klein R. Mitogen-inducible gene 6 is an endogenous inhibitor of HGF/Met-induced cell migration and neurite growth. The Journal of cell biology 171: 337–348, 2005.

28. Portero-Otin M, Pamplona R, Bellmunt MJ, Ruiz MC, Prat J, Salvayre R, and Negre-Salvayre A. Advanced glycation end product precursors impair epidermal growth factor receptor signaling. Diabetes 51: 1535–1542, 2002.

29. Segatto O, Anastasi S, and Alema S. Regulation of epidermal growth factor receptor signalling by inducible feedback inhibitors. Journal of cell science 124: 1785–1793, 2011.

30. Srinivasan S, Ohsugi M, Liu Z, Fatrai S, Bernal-Mizrachi E, and Permutt MA. Endoplasmic reticulum stress-induced apoptosis is partly mediated by reduced insulin signaling through phosphatidylinositol 3-kinase/Akt and increased glycogen synthase kinase-3beta in mouse insulinoma cells. Diabetes 54: 968–975, 2005.

31. Tran VV, Chen G, Newgard CB, and Hohmeier HE. Discrete and complementary mechanisms of protection of beta-cells against cytokine-induced and oxidative damage achieved by bcl-2 overexpression and a cytokine selection strategy. Diabetes 52: 1423–1432, 2003.

32. Wrede CE, Dickson LM, Lingohr MK, Briaud I, and Rhodes CJ. Protein kinase B/Akt prevents fatty acid-induced apoptosis in pancreatic beta-cells (INS-1). The Journal of biological chemistry 277: 49676–49684, 2002.

33. Xu D, Makkinje A, and Kyriakis JM. Gene 33 is an endogenous inhibitor of epidermal growth factor (EGF) receptor signaling and mediates dexamethasone-induced suppression of EGF function. The Journal of biological chemistry 280: 2924–2933, 2005.

34. Zarrouki B, Benterki I, Fontes G, Peyot ML, Seda O, Prentki M, and Poitout V. Epidermal Growth Factor Receptor Signaling Promotes Pancreatic beta-Cell Proliferation in Response to Nutrient Excess in Rats through mTOR and FOXM1. Diabetes 2013.

